# MASIv2 enables standardization and integration of multi-modal single-cell and spatial omics data with one general framework

**DOI:** 10.1101/2023.05.15.540808

**Authors:** Yang Xu, Sikander Hayat

**Affiliations:** Data Sciences Platform, Broad Institute of Harvard and MIT, Cambridge, USA; Institute of Experimental Medicine and Systems Biology, Uniklinik RWTH Aachen, Aachen, Germany

## Abstract

Data annotation and integration are two common tasks in large-scale and collaborative single-cell research. Rapid technological advancements have made diverse single-cell and spatial data modalities available. This data deluge brought up great challenges in data annotation and integration. Though different biological modalities preserve shared features to define the same cellular system, they often present unique angles to unravel a multi-level understanding about this system. Here, we present one general framework that uses modality-shared and -specific features for annotation and integration of single-cell and spatial omics data. We benchmark our framework with existing methods across different datasets and demonstrate its application in two real world tasks.

## 1. Introduction

### 1.1. The fundamental task in single-cell data analysis

Cell-type annotation is an essential task needed for most downstream single-cell analyses. Single-cell transcriptome (scRNA-seq), the predominant technology in single-cell research, has generated numerous large-scale datasets. Manual annotation for these datasets becomes time-consuming and less reproducible. Thus, there is a great demand to set up trustworthy computational standards. Fortunately, many tools, either based on classification or known markers, were proposed to address automated annotation of scRNA-seq data. Recently, the quantity of other single-cell biological modalities has readily increased. However, existing tools for annotating scRNA-seq data may not be capable of handling other biological modalities. Tools for multi-modal annotation are also in a great demand.

### 1.2. Principles of multi-modal data integration

To deal with multi-modal annotation, we cannot avoid integrating different data modalities first. Multi-modal data integration can be summarized into 3 categories: vertical, horizontal, and diagonal integration (Argelaguet et al., 2021). Horizontal integration requires shared features to anchor data from different modalities, while shared cells serve the same purpose in vertical integration. Distinctly, diagonal integration does not require shared features or cells for anchoring data. In a follow-up comment, we argued that diagonal integration is not a safe practice, leaving horizontal and vertical integration as two viable options (Xu & McCord, 2022).

### 1.3. Need for a simple solution

For both cell-type annotation and data integration, we observe a trend towards big and resource-intensive models. These computationally expensive methods would potentially limit the single-cell scientific research to resource-rich laboratories and exclude researchers with fewer resources to participate in large-scale collaborative projects. Moreover, big models do not necessarily return biologically meaningful outcomes. Evidence shows that complicated methods are not necessarily better at preserving the underlying biological information (Luecken et al., 2022). In line with that, we propose a simple yet effective computational framework to handle annotation and integration of multi-modal single-cell and spatial omics data.

## 2. Related Work

### 2.1. A simple method can be as good as sophisticated ones

Two benchmark studies demonstrated that sophisticated models do not outperform a simple linear model for the task of cell-type annotation for scRNA-seq data (Abdelaal et al., 2019; Köhler et al., 2019). Additionally, (Pliner et al., 2019) and (Domínguez Conde et al., 2022) proposed logistic regression based methods for cell-type annotation, and both methods could achieve great accuracies in this task. Independently, we used putative cell-type markers to handle different cases of scRNA-seq data annotation (Xu et al., 2022a; 2023). All these studies demonstrate that a simple solution can be satisfactory for the problem of cell-type annotation for scRNA-seq data.

### 2.2. Success of horizontal integration

Though we argue that cell-type annotation for scRNA-seq data is a simple task, a well-recognized practice to label other modalities of single-cell and spatial omics data is still missing. One solution is transferring labels from scRNA-seq data to other modalities, but this task will be coupled with multi-modal integration. Fortunately, numerous experimental studies confirm that feature correlations exist among different biological modalities. For instance, highly expressed *VCAN* gene could be used to define CD14+ monocytes in scRNA-seq data, while open chromatin in the *VCAN* gene body serves the same purpose in single-cell chromatin accessibility (scATAC-seq) data. This kind of feature correlations do not only exist between scRNA-seq and scATAC-seq data, but also between many modalities, including DNA methylation (Lee et al., 2019), histone modification (Zhu et al., 2021), and chromatin structure (Shen et al., 2022). Because of the apparent feature correlations across different modalities, horizontal integration that anchors shared features thrived as a major solution to multi-modal integration. As the growing availability of multi-modal single-cell data, some horizontal methods such as Seurat (Butler et al., 2018), Harmony (Korsunsky et al., 2019), and LIGER (Gao et al., 2021) that rely on large matrix manipulation, became less capable of handling multi-modal data integration efficiently. Neural-network-based integration methods, then, rise as alternatives. These methods were shown to handle large-scale single-cell integration more efficiently, because of the manner of handling data into mini batches (Lin et al., 2022; Xiong et al., 2022; Xu et al., 2022b).

### 2.3. The use of graph in single-cell data analysis

All those horizontal integration methods above share one thing in common: they exhaustively take advantage of shared features for the purpose of data integration, regardless of the inherent modality-specific differences. As different biological modalities present the same cellular system in different views, modality-specific information is critical to a comprehensive understanding of these systems. (Jain et al., 2021) proposed a method named MultiMAP, which does not discard modality-specific information during horizontal integration. MultiMAP builds graphs with modality-specific features, and it integrates graphs by finding an optimal alignment with shared features. Aligning two graphs can be computationally expensive to solve. Nowadays, single-cell omics data are tremendously large, and the size of integrated atlases can easily be in the range of millions of cells. Therefore, building graphs with millions of nodes in each modality and aligning them can be challenging.

## 3. Proposed Framework

Previously, we developed MASI after extensively testing the ability of putative cell-type markers for annotation and integration of scRNA-seq data (Xu et al.. 2022a; 2023). Built upon our previous studies, we aim to extend the marker-assisted annotation and integration strategy to multi-modal single-cell and spatial omics data. Keeping our commitment to simplicity and inclusion, the new MASIv2 is still computationally inexpensive and efficient, but it expands its capacity to cover additional modalities of single-cell, and spatial omics data. Briefly, we set up our task as annotating N cells of each modality into K cell types with M known markers of each cell type and further integrating Q modalities all at once (Figure 1).

**Figure 1.**
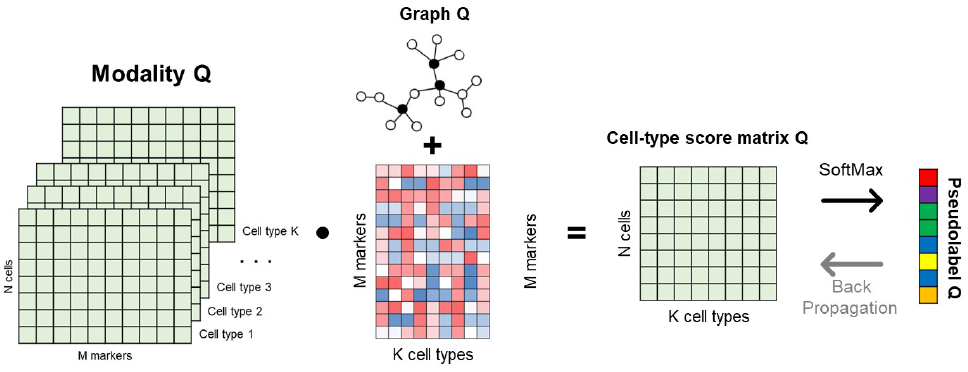
The general framework for annotating and integrating multi-modal single-cell and spatial omics data. The N by M by K matrix of each modality was built with marker genes as shared features. Each graph was constructed with modality-specific features. The N by M by K matrix is transformed by the trainable M by K weight matrix after passing through modality-specific graph. SoftMax was applied to the N by K output matrix to predict K cell types.

### 3.1. Encode prior knowledge as trainable weights

Our proposed model in MASIv2 transforms the shared gene feature space *X*_*q*_ to cell-type feature space *z*_*q*_, where q stands for one specific modality. For each modality, we aim to learn a unique transformation that maps *N*_*q*_ * *K*_*q*_ * *M*_*q*_ input into *N*_*q*_ * *K*_*q*_ output matrix. For all Q modalities, we need *K*_*q*_ * *M*_*q*_ * *Q* trainable weights. For example, the multi-modal mouse primary motor cortex data containing 24 cell types, with 30 identified markers per cell type, for 4 biological modalities, would require only 24*30*4 = 2880 weishts.

### 3.2. Represent modalities as graphs

Then, we construct graphs for each modality using modality-specific features. The modality-specific graph preserves the distinct structure of that modality and will be specifically used to update *K*_*q*_ * *M*_*q*_ weights. However, graph computation is usually expensive. Incorporating graph could add substantial computation burden and further prohibit users with few computational resources to use our framework. After surveying different architectures of graphs, we avoided using any architectures that have numerous trainable weights. In fact, linear graph model can achieve state-of-the-art performance, promising a computation-light solution for graph representation (Thekumparampil et al., 2018; Wu et al., 2019). Here, we use attention-based graph layer to incorporate modality-specific graphs into our framework (Thekumparampil et al., 2018). As the attention-based graph layer does not have any weights to train, this makes graph computation on a personal laptop feasible. Meanwhile, building graphs with over 1 million nodes is still formidable if users do not have high-end computation resources. Our proposal here is to divide the whole dataset into multiple small chunks and build sub-graph for each chunk. A similar solution in our previous study has showed optimal performance over multiple single-cell datasets (Xu et al., 2023).

### 3.3. A bad teacher, but a good student

We pass the shared gene features *X*_*q*_ through modality-specific attention-based graphs before we transform them into cell-type feature *z*_*q*_ with the *K*_*q*_ * *M*_*q*_ weights. The output is a *N*_*q*_ * *K*_*q*_ matrix for each modality. Applying SotfMax, we could assign *K*_*q*_ cell types to these *N*_*q*_ cells. We initialize all *K*_*q*_ \**M*_*q*_ trainable weights to 1. The initialized model can annotate single-cell or spatial omics data, but it is likely that we will end up with erroneous labels. Even so, this bad model could serve as a teacher to generate pseudo labels. Then, pseudo labels can help the student model update weights. Eventually, the student model would correct mislabeling done by its teacher. (Caron et al., 2018) showed that stochastic learning has the ability to learn from an erroneous start and evolves to amplify correct signals. Self-supervised learning that relies on self-labeling supports this idea and demonstrates a certain degree of capacity of correctness (Asano et al., 2019). To greatly improve student from its teacher, we apply knowledge distillation, in which we introduce temperature term *π*_1_ (>= 10) into *softmax*(*z*_*q*_*/π*_1_) to generate smooth probabilities (Hinton et al., 2015).

### 3.4. A smooth SoftMax, and a sharp SoftMax

Use of knowledge distillation presents one side of SoftMax - smoothness. The ability to correct errors made by teacher would saturate at one point, as smoothness does not discourage model to generate labels with high confidence. Inspired by the opposite use of temperature in contrastive learning, where lower temperature is used to generate sharp Soft-Max outputs to learn discriminative representation (Chen et al., 2020). Here, we introduce annotated scRNA-seq data as reference. We did not use the annotated scRNA-seq data in training directly, for the sake of efficiency. Instead, we averaged the shared gene features *X* of the annotated scRNA-seq data by cell-type label to get *X*_c_ of cell-type centroids. Each centroid is a *K* * *M* length vector. Then we introduce the second temperature term *π*_2_ (<= 0.1) into softmax(z_q_/*π*_2_) to generate sharp probabilities *p*_*q*_. Next, we apply Mean squared error loss to reduce the difference between (*K*_*q*_ * *M*_*q*_) * *X*_*q*_ and *p*_*q*_ • *X*_*c*_. We can skip this part if an annotated scRNA-seq data is not available.

### 3.5. Linear correction for data integration

The difficulty of aligning latent representations of different modalities is that each latent dimension can be arbitrary in different modalities. MASIv2 avoids this problem by projecting gene feature space *X*_*q*_ to cell-type feature space *z*_*q*_. Thus, modalities are “pre-aligned” in the cell-type feature space *z*_*q*_. The remaining problem of modality alignment can be solved by linear correction. For simplicity, we anchor one modality as the source and map all other modalities to the source. Use of graph in previous steps gives us the advantage of identifying key points from each modality specific graph. Unlike MultiMAP that directly finds the best alignment across the whole graph, we only search for a solution with key points. Here, we use Louvain community detection to identify a group of key points for each modality. Then, we sample key points for each cell type from different modailties as pairs. Next, we train a linear regression model that can match two key points of the same cell type but from two modalities. This linear correction only adds *K* * (Q −1) trainable weights with Q − 1 bias terms into our proposed framework. Take the multi-modal mouse primary motor cortex as example, we need to further introduce (24 + 1) * (4 + 1) = 75 trainable weights for purpose of modality alignment. Combining trainable weights introduced earlier, the final model for multi-modal mouse primary motor cortex data would have 2955 weights in total. Compared to any deep-learning based integration models, our framework is still substantially small.

## 4. Experiments

### 4.1. Benchmarking

We benchmarked MASIv2 with 4 tools for the task of cross-modal label transferring. We collected 4 scATAC-seq for annotation and 4 scRNA-seq datasets from the same 4 tissues as reference (Miao et al., 2021; Kuppe et al.. 2021; Ma et al., 2020; Granja et al., 2019). Cell-type labels reported in the original studies served as the ground truth for evaluation. As annotation resolutions and naming styles between the scATAC-seq and its corresponding scRNA-seq data do not always match, we used Adjusted rand index (ARI) for evaluation. First, we observed that the student model of MASIv2 had substantial improvement from its teacher model, indicating a good power of label correction during learning (Table 1). Second, MASIv2 had the highest ARI scores across 3 datasets, except mouse skin data (Table 1). These two results show that MASIv2 is competent against existing methods in cross-modal annotation.

**Table 1.** Cross-modal annotation accuracies (ARI) of 5 computational methods in 4 scATAC-seq datasets.

| METHOD | MOUSE KIDNEY | HUMAN KIDNEY | MOUSE SKIN | HUMAN PBMC |
| --- | --- | --- | --- | --- |
| SEURAT | 0.653 | 0.707 | **0.564** | 0.592 |
| HARMONY | 0.483 | 0.628 | 0.519 | 0.587 |
| LIGER | 0.532 | 0.567 | 0.181 | 0.471 |
| SCICAN | 0.448 | 0.660 | 0.527 | 0.585 |
| MASIV2 (TEACHER) | 0.393 | 0.682 | 0.201 | 0.294 |
| MASIV2 (STUDENT) | **0.760** | **0.722** | 0.532 | **0.620** |

### 4.2. Application in spatial omics data

In spatial omics data, each spot in space is surrounded by other spots, and all these spots form a natural graph. Because this graph is constructed based on how close they are in space, it is possible that spots can form edges even though they may not have any similarities. At those border regions, it is more difficult to assign cell-type labels to those spots. A possible solution is to give users choices on how much cell-type identity of each spot would be influenced by its neighboring spots. To do that, we introduced a third temperature parameter *π*_3_ which was applied in the attention mechanism of the graph layer to control sharpness of the SoftMax output. A sharper SoftMax discounts attention scores contributed by dissimilar neighboring spots and gives higher attention to similar neighboring spots. We set *π*_3_ as 0.1 for all cases of spatial omics data analysis. (Marshall et al., 2022) applied Slide-seqV2 to study autosomal dominant polycystic kidney disease (ADPKD) in mouse. We applied MASIv2 to this Slide-seqV2 data, and MASIv2 revealed distinct spatial patterns between control and ADPKD (Figure 2). Meanwhile, the cell-type feature *z* returned by MASIv2 could also be interpreted as deconvolution of cell types. This paints a more clear picture about spatial pattern in mouse ADPKD. For example a denser aggregation of proximal straight tubule (PST) and presence of macrophage (Macro) in ADPKD samples (Figure 2).

**Figure 2.**
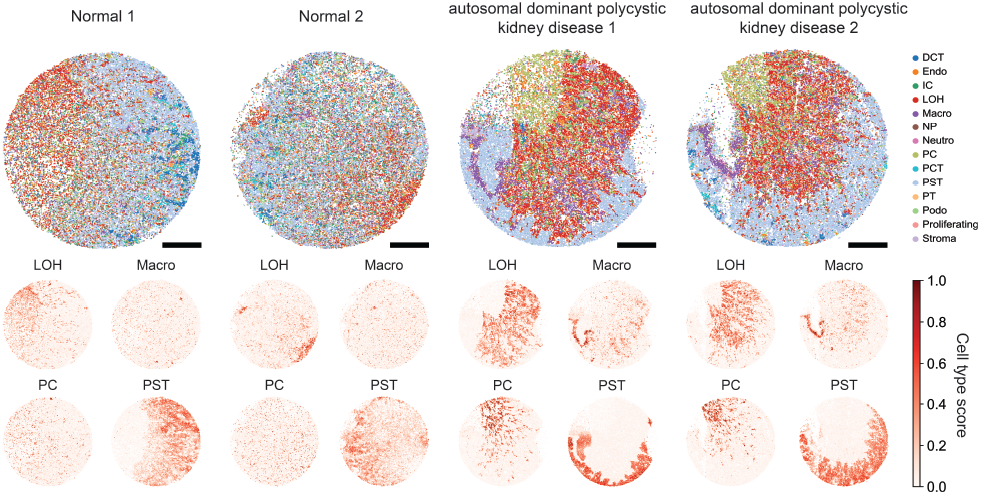
Annotation of spatial mouse kidney data. Two control tissues and two samples with ADPKD are presented. Cell type scores *softmax* (*z*_*q*_) predicted by MASIv2 are shown to highlight distinct cellular difference between control and disease samples.

### 4.3. Integrating all modalities at once

Finally, we demonstrate that MASIv2 can scale to integrate multiple modalities at once, including gene expression, chromatin accessibility, DNA methylation, and chromatin structure (BICCN, 2021; Tan et al., 2021). We previously identified marker genes for cell types in mouse primary cortex (Xu et al., 2023). The identified marker genes serve to build *K*_*q*_ \**M*_*q*_ for each modality. As no suitable reference data was available, we first annotated gene expression modality without the step mentioned in section **3.4**. Then, the annotated gene expression modality was used as reference to help train models for the other 3 modalities. Annotations by MASIv2 have good agreement with author-verified annotation, even chromatin structure (Figure 3).

**Figure 3.**
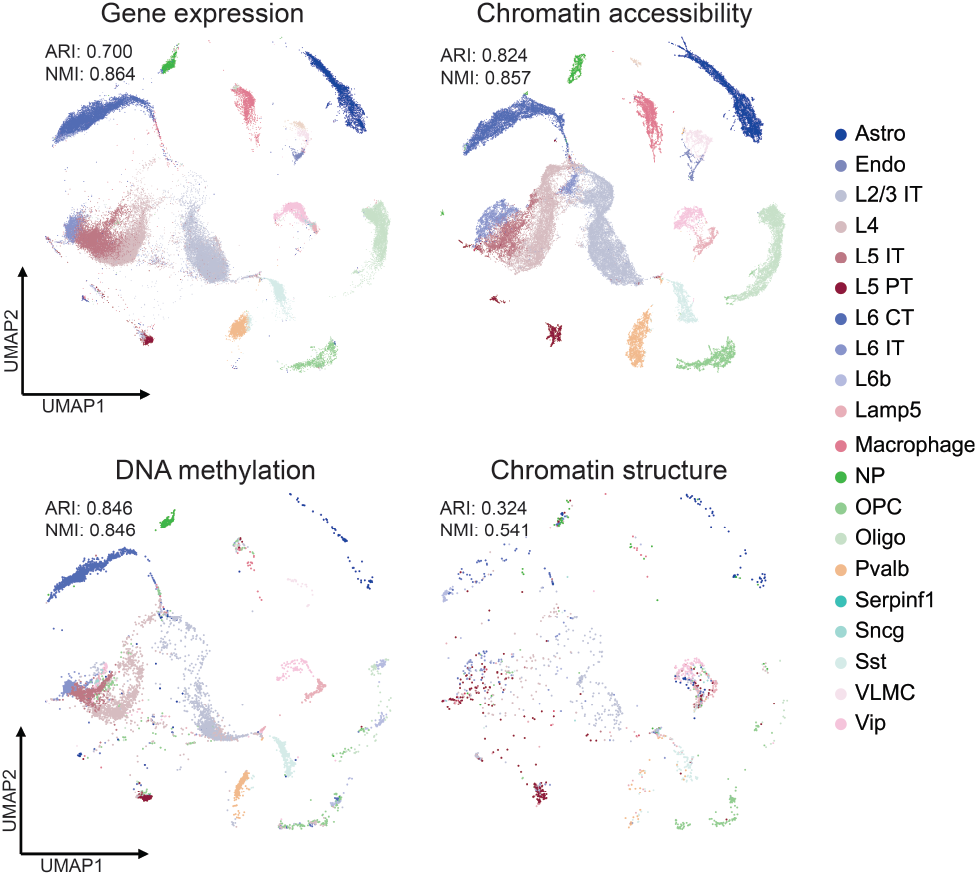
Integrated mouse motor cortex data across 4 modalities, including gene expression, chromatin accessibility, DNA methylation, and chromatin structure. Annotation accuracy is measured via ARI and NMI against author-verified cell type labels.

## 5. Conclusion

We show the utility of small computational models in single-cell research. Going small as an alternative could still be effective as big models in some cases for biological problems. In this study, we demonstrate MASIv2, a general framework with a small set of weights, can achieve good accuracy of cross-modal annotation, while effectively integrating multi-modal single-cell and spatial omics data.

## 6. Funding

SH is supported in part by Leducq Seed Award and RWTH START Grant.

## 7. Disclosures

SH reports grants from Novo Nordisk and Askbio GmbH.

## Notes

### Competing Interest Statement

The authors have declared no competing interest.

## References

Abdelaal, T., Michielsen, L., Cats, D., Hoogduin, D., Mei, H., Reinders, M. J., and Mahfouz, A. A comparison of automatic cell identification methods for single-cell rna sequencing data. Genome biology, 20:1–19, 2019.

Argelaguet, R., Cuomo, A. S., Stegle, O., and Marioni, J. C. Computational principles and challenges in single-cell data integration. Nature biotechnology, 39(10):1202–1215, 2021.

Asano, Y. M., Rupprecht, C., and Vedaldi, A. Self-labelling via simultaneous clustering and representation learning. arXiv preprint 1911.05371, 2019.

BICCN. A multimodal cell census and atlas of the mammalian primary motor cortex. Nature, 598(7879):86–102, 2021.

Butler, A., Hoffman, P., Smibert, P., Papalexi, E., and Satija, R. Integrating single-cell transcriptomic data across different conditions, technologies, and species. Nature biotechnology, 36(5):411–420, 2018.

Caron, M., Bojanowski, P., Joulin, A., and Douze, M. Deep clustering for unsupervised learning of visual features. In Proceedings of the European conference on computer vision (ECCV), pp. 132–149, 2018.

Chen, T., Kornblith, S., Norouzi, M., and Hinton, G. A simple framework for contrastive learning of visual representations. In International conference on machine learning, pp. 1597–1607. PMLR, 2020.

Domínguez Conde, C., Xu, C., Jarvis, L., Rainbow, D., Wells, S., Gomes, T., Howlett, S., Suchanek, O., Polanski, K., King, H., et al. Cross-tissue immune cell analysis reveals tissue-specific features in humans. Science, 376 (6594):eabl5197, 2022.

Gao, C., Liu, J., Kriebel, A. R., Preissl, S., Luo, C., Castanon, R., Sandoval, J., Rivkin, A., Nery, J. R., Behrens, M. M., et al. Iterative single-cell multi-omic integration using online learning. Nature biotechnology, 39(8): 1000–1007, 2021.

Granja, J. M., Klemm, S., McGinnis, L. M., Kathiria, A. S., Mezger, A., Corces, M. R., Parks, B., Gars, E., Liedtke, M., Zheng, G. X., et al. Single-cell multiomic analysis identifies regulatory programs in mixed-phenotype acute leukemia. Nature biotechnology, 37(12):1458–1465, 2019.

Hinton, G., Vinyals, O., and Dean, J. Distilling the knowledge in a neural network. arXiv preprint 1503.02531, 2015.

Jain, M. S., Polanski, K., Conde, C. D., Chen, X., Park, J., Mamanova, L., Knights, A., Botting, R. A., Stephenson, E., Haniffa, M., et al. Multimap: dimensionality reduction and integration of multimodal data. Genome biology, 22 (1):1–26, 2021.

Köhler, N. D., Büttner, M., and Theis, F. J. Deep learning does not outperform classical machine learning for celltype annotation. BioRxiv, pp. 653907, 2019.

Korsunsky, I., Millard, N., Fan, J., Slowikowski, K., Zhang, F., Wei, K., Baglaenko, Y., Brenner, M., Loh, P.-r., and Raychaudhuri, S. Fast, sensitive and accurate integration of single-cell data with harmony. Nature methods, 16(12): 1289–1296, 2019.

Kuppe, C., Ibrahim, M. M., Kranz, J., Zhang, X., Ziegler, S., Perales-Patón, J., Jansen, J., Reimer, K. C., Smith, J. R., Dobie, R., et al. Decoding myofibroblast origins in human kidney fibrosis. Nature, 589(7841):281–286, 2021.

Lee, D.-S., Luo, C., Zhou, J., Chandran, S., Rivkin, A., Bartlett, A., Nery, J. R., Fitzpatrick, C., O’Connor, C., Dixon, J. R., et al. Simultaneous profiling of 3d genome structure and dna methylation in single human cells. Nature methods, 16(10):999–1006, 2019.

Lin, Y., Wu, T.-Y., Wan, S., Yang, J. Y., Wong, W. H., and Wang, Y. R. scjoint integrates atlas-scale single-cell rna-seq and atac-seq data with transfer learning. Nature biotechnology, 40(5):703–710, 2022.

Luecken, M. D., Büttner, M., Chaichoompu, K., Danese, A., Interlandi, M., Müller, M. F., Strobl, D. C., Zappia, L., Dugas, M., Colomé-Tatché, M., et al. Benchmarking atlas-level data integration in single-cell genomics. Nature methods, 19(1):41–50, 2022.

Ma, S., Zhang, B., LaFave, L. M., Earl, A. S., Chiang, Z., Hu, Y., Ding, J., Brack, A., Kartha, V. K., Tay, T., et al. Chromatin potential identified by shared single-cell profiling of rna and chromatin. Cell, 183(4):1103–1116, 2020.

Marshall, J. L., Noel, T., Wang, Q. S., Chen, H., Murray, E., Subramanian, A., Vernon, K. A., Bazua-Valenti, S., Liguori, K., Keller, K., et al. High-resolution slide-seqv2 spatial transcriptomics enables discovery of disease-specific cell neighborhoods and pathways. Iscience, 25 (4):104097, 2022.

Miao, Z., Balzer, M. S., Ma, Z., Liu, H., Wu, J., Shrestha, R., Aranyi, T., Kwan, A., Kondo, A., Pontoglio, M., et al. Single cell regulatory landscape of the mouse kidney highlights cellular differentiation programs and disease targets. Nature communications, 12(1):2277, 2021.

Pliner, H. A., Shendure, J., and Trapnell, C. Supervised classification enables rapid annotation of cell atlases. Nature methods, 16(10):983–986, 2019.

Shen, S., Zheng, Y., and Keleş, S. scgad: single-cell gene associating domain scores for exploratory analysis of schi-c data. Bioinformatics, 38(14):3642–3644, 2022.

Tan, L., Ma, W., Wu, H., Zheng, Y., Xing, D., Chen, R., Li, X., Daley, N., Deisseroth, K., and Xie, X. S. Changes in genome architecture and transcriptional dynamics progress independently of sensory experience during post-natal brain development. Cell, 184(3):741–758, 2021.

Thekumparampil, K. K., Wang, C., Oh, S., and Li, L.-J. Attention-based graph neural network for semisupervised learning. arXiv preprint 1803.03735, 2018.

Wu, F., Souza, A., Zhang, T., Fifty, C., Yu, T., and Weinberger, K. Simplifying graph convolutional networks. In International conference on machine learning, pp. 6861–6871. PMLR, 2019.

Xiong, L., Tian, K., Li, Y., Ning, W., Gao, X., and Zhang, Q. C. Online single-cell data integration through projecting heterogeneous datasets into a common cell-embedding space. Nature Communications, 13(1):6118, 2022.

Xu, Y. and McCord, R. P. Diagonal integration of multimodal single-cell data: potential pitfalls and paths forward. Nature Communications, 13(1):3505, 2022.

Xu, Y., Baumgart, S. J., Stegmann, C. M., and Hayat, S. Maca: marker-based automatic cell-type annotation for single-cell expression data. Bioinformatics, 38(6):1756–1760, 2022a.

Xu, Y., Begoli, E., and McCord, R. P. scican: single-cell chromatin accessibility and gene expression data integration via cycle-consistent adversarial network. npj Systems Biology and Applications, 8(1):33, 2022b.

Xu, Y., Kramann, R., McCord, R. P., and Hayat, S. Masi enables fast model-free standardization and integration of single-cell transcriptomics data. Communications Biology, 6(1):465, 2023.

Zhu, C., Zhang, Y., Li, Y. E., Lucero, J., Behrens, M. M., and Ren, B. Joint profiling of histone modifications and transcriptome in single cells from mouse brain. Nature methods, 18(3):283–292, 2021.

